# Optic radiations representing different eccentricities age differently

**DOI:** 10.1101/2022.10.03.510705

**Authors:** John Kruper, Noah C. Benson, Sendy Caffarra, Julia Owen, Yue Wu, Aaron Y. Lee, Cecilia S. Lee, Jason Yeatman, Ariel Rokem

## Abstract

The white matter pathways that carry information from the foveal, macular, and peripheral visual fields have distinct biological properties. The optic radiations (OR) carry foveal and peripheral information from the thalamus to the primary visual cortex (V1) through adjacent but separate pathways in the white matter. Here, we perform white matter tractometry using pyAFQ ^1^ on a large sample of diffusion MRI (dMRI) data from the UK Biobank dataset (UKBB; N=6021; age 45-81). We use pyAFQ to characterize white matter tissue properties in parts of the OR that transmit information about the foveal, macular, and peripheral visual fields, and to characterize the changes in these tissue properties with age. We find that (1) independent of age there is higher fractional anisotropy, lower mean diffusivity, and higher mean kurtosis in the foveal and macular OR than in peripheral OR, consistent with denser, more organized nerve fiber populations in foveal/parafoveal pathways, and (2) age is associated with increased diffusivity and decreased anisotropy and kurtosis, consistent with decreased density and tissue organization with adulthood aging. However, anisotropy in foveal OR decreases faster with age than in peripheral OR, while diffusivity increases faster in peripheral OR, suggesting foveal/peri-foveal OR and peripheral OR differ in how they age.

## Introduction

Visual perception of peripheral and foveal eccentricities differs substantially, suggesting qualitatively different computational mechanisms governing each of these parts of the visual field. These differences in perception and computation stem from structural and functional differences between foveal and peripheral representations at every stage of the visual system. In the retina, the fovea occupies a privileged position with no occlusion by blood vessels or incident axons, and a one-to-one ratio of receptors to ganglion cells ^2^. The magnification of the fovea in the retina is further enhanced as information travels to the cortex, with approximately 50% of the primary visual cortex (V1) devoted to the central 12 degrees of vision ^3^. Nearby points in the visual field are represented by neighboring points on the surface of the visual cortex ^4^. This mapping is a consequence of the ordered set of projections from the retina to the lateral geniculate nucleus (LGN) and from the LGN to the V1. The projections from LGN to V1 through the optic radiations (OR), in particular, are known to contain a retinotopic organization, with nerve fibers transmitting information about neighboring parts of the visual field traveling close to each other. Information about nearby points in the visual field travel through adjacent bundles of axons in the white matter, as reflected in the fact that damage to the OR results in scotomas in predictable parts of the visual field ^5^. Here, we asked whether there are systematic differences between the white matter projections that transmit information about the central visual field and more peripheral eccentricities.

New large and openly available datasets like the UK Biobank ^6^ (UKBB) provide an opportunity to study within and cross-individual differences in the physical properties of the OR at unprecedented scale. Here, we used a large sample (n>6,000) from the UKBB dataset to ask whether there are differences in the tissue properties of parts of the OR that contain the axons that transmit information about different parts of the visual field. Furthermore, aging is reflected in measurable changes in the physical properties of the tissue ^7,8^. In the retina, the fovea has a different trajectory of aging than the periphery ^9,10^. The large range of ages represented in the UKBB sample also allows us to ask whether distinct effects of aging can be measured in different parts of the OR.

To assess white matter tissue properties, we analyzed UKBB diffusion-weighted MRI (dMRI), which measures the random motion of water within brain tissue ^11^. The voxels of dMRI measurements are typically on the order of 1mm in size, but dMRI measurements are sensitive to microstructural features at the scale of microns. In the white matter of the brain, where axon bundles are densely packed, water more readily diffuses along the length of the axons. In voxels containing white matter, diffusion measurements are anisotropic and indicate the directions of fiber bundles in these voxels. Computational tractography uses this directional diffusion information to generate estimates of the trajectories of white matter pathways between different parts of the brain. In the white matter between the LGN and V1, dMRI can be used to accurately delineate the trajectory of the optic radiations (OR) ^12–14^. Furthermore, owing to the systematic mapping of the visual field in the visual cortex, parts of the OR that transmit information about different parts of the visual field can be systematically parsed based on their endpoints close to the visual cortex ^12,15^.

In addition to estimating macrostructural white matter tracts, diffusion MRI data can also be used to assess the microstructural properties of the white matter tissue. Here, we used the diffusional kurtosis model (DKI) ^16,17^. DKI extends the classical diffusion tensor model (DTI ^18^) to account for non-Gaussian diffusion that occurs in heterogeneous white matter tissue and due to restriction of water diffusion within densely packed axons and other cellular structures. The degree of non-Gaussianity is quantified via the mean kurtosis (MK). In addition, we also used DTI metrics: the average diffusion across directions (mean diffusivity or MD), and the fractional anisotropy (FA). FA is a normalized variance of the diffusion across directions, which is bounded between 0 (isotropic diffusion) and 1 (anisotropic diffusion). DTI-derived parameters, such as FA and MD are highly sensitive to biological change and to differences between individuals, but unfortunately, they are also non-specific. For example, FA tends to decrease with demyelination ^19^, leading some to interpret this parameter as indicative of “white matter integrity”. However, it also decreases in voxels in which more than one major fiber population is present, suggesting that caution should be taken in this interpretation ^20^. MK helps reduce the ambiguity and further constrains the interpretation of FA and MD. For example, both damage to a population of fibers and the addition of crossing fibers reduce FA, but the former causes a decrease in MK, while the latter would cause an increase in MK ^17^. Tissue properties calculated with DKI also relate to microstructural changes present in brain diseases ^21,22^.

To assess the tissue properties in different sub-bundles of the OR, corresponding to different parts of the visual field, and their change with age, we used pyAFQ (https://yeatmanlab.github.io/pyAFQ; ^1^), an open-source software pipeline that implements tractometry based on the Automated Fiber Quantification approach ^23^. In this approach, white matter pathways, such as the OR, are automatically identified based on anatomical landmarks, and diffusion properties are quantified along the trajectory of the bundle. Tractometry is used to characterize the physical properties of the major white matter pathways along their length, taking into account systematic variability that exists in these properties throughout the length of the major white matter pathways. We recently demonstrated that this process is both reliable in test-retest data, as well as robust to variations in computational methodology ^1^, despite variability in the results of tractography ^24^. Using this approach, we demonstrate a differentiation between white matter properties of different sub-bundles of the OR, and show that age-related changes in tissue properties systematically differ between the sub-bundles. These findings suggest different aging trajectories for different populations of nerve fibers even within the same anatomical pathway.

## Methods

### Subjects

We selected dMRI data from 7,438 subjects that had both diffusion MRI data and OCT measurements in each eye ^6,25^. Of these, we selected 6,021 subjects that are classified as having no eye problems/disorders based on an ACE touchscreen question: “Has a doctor told you that you have any of the following problems with your eyes? (You can select more than one answer)” (see UKBB data field 6148). Population characteristics for this sample are shown in Figure 1.

**Figure 1:**
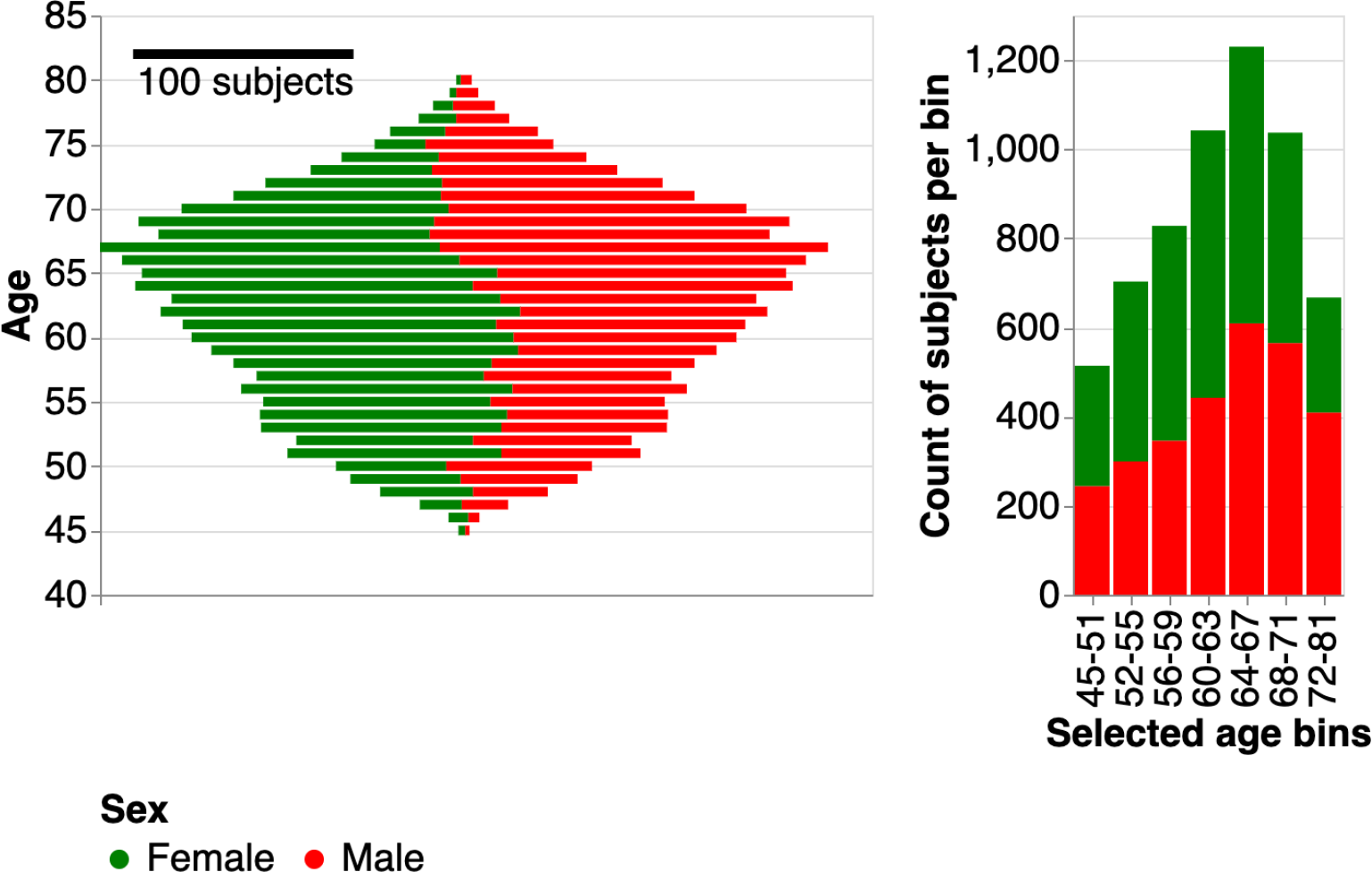
Population characteristics of UKBB subjects. In each panel, gender is denoted by color. In the left panel, we plot the age distribution. Note that younger subjects are majority female, while older subjects are majority male. In the right panel, we break down the subjects into age bins. The middle age bins all span 4 years. The first and last age bins were selected to have similar numbers of subjects in each bin to the middle age bins. These age bins are used to group subjects and to visualize changes in tract profiles with age.

### AFQ configuration

#### Data

We used preprocessed diffusion MRI data that were processed and released by the UK Biobank team. The acquisition protocol has been described elsewhere ^27^, and we provide here only some details. Data were acquired with a spatial resolution of 2-by-2-by-2 mm^3^. TE/TR = 14.92/3600 msec. Five volumes were acquired with no diffusion weighting (b=0), and 50 volumes were acquired for each of two diffusion weightings: b=1,000 s/mm^2^ and b=2,000 s/mm^2^. In addition to these 105 volumes acquired with an anterior-to-posterior phase encoding direction, and additional 6 b=0 volumes were acquired with posterior-to-anterior phase encoding direction and subsequently used for EPI distortion correction. Preprocessing was also described elsewhere ^25^, and we provide only some details. Briefly, head motion and eddy currents were corrected using the FSL “eddy” software, including correction of outlier slices. Subsequently gradient distortion correction was performed. Non-linear registration using FNIRT was used to map the individual-level data to the MNI template.

#### Tractography and Registration

Residual bootstrap tractography ^28^ was used to delineate the trajectory of optic radiations in each subject’s individual data. We used a GPU-accelerated implementation of this method ^28^, limiting tractography to the posterior half of the brain. Based on the known trajectory of the optic radiations, we defined inclusion, exclusion, and endpoint regions of interest (ROIs) within the core white matter in each hemisphere. These were registered to each subject’s anatomy using the FNIRT non-linear warp and 64 seeds were uniformly distributed in each voxel of each ROI.

#### Bundle Recognition

Bundle recognition used the pyAFQ software (https://github.com/yeatmanlab/pyAFQ; ^1^), which implements a procedure very similar to the one described by Yeatman *et al*.^23^. The software finds streamlines that belong to a white matter pathway by defining waypoint ROIs within the core of the white matter along the trajectory of the pathway. The software allows additional criteria for inclusion or exclusion of tractography streamlines; we used the following criteria to recognize the OR: streamlines (1) do not pass through the sagittal midline of the brain; (2) have at least one point that is within 3 mm of both of the inclusion ROIs; (3) do not have any point that is within 3 mm from the exclusion ROI; (4) terminate within 3 mm of the two endpoint ROIs (one in the thalamus and the other in V1) ^14^. After defining this group of streamlines an additional cleaning procedure was applied to remove streamlines that were outliers in terms of their length and trajectory. Subsequently, streamlines were divided into foveal OR (fOR), macular OR (mOR), or peripheral OR (pOR) based on the anatomical position of their termination in V1, using eccentricity from the retinotopic prior of Benson and Winawer ^29^ and masked with the V1 location in the AICHA atlas ^30^ (Figure 2). The eccentricity ranges were: fOR, <=3°; mOR, >3°, <=7°; pOR, >7°.

**Figure 2:**
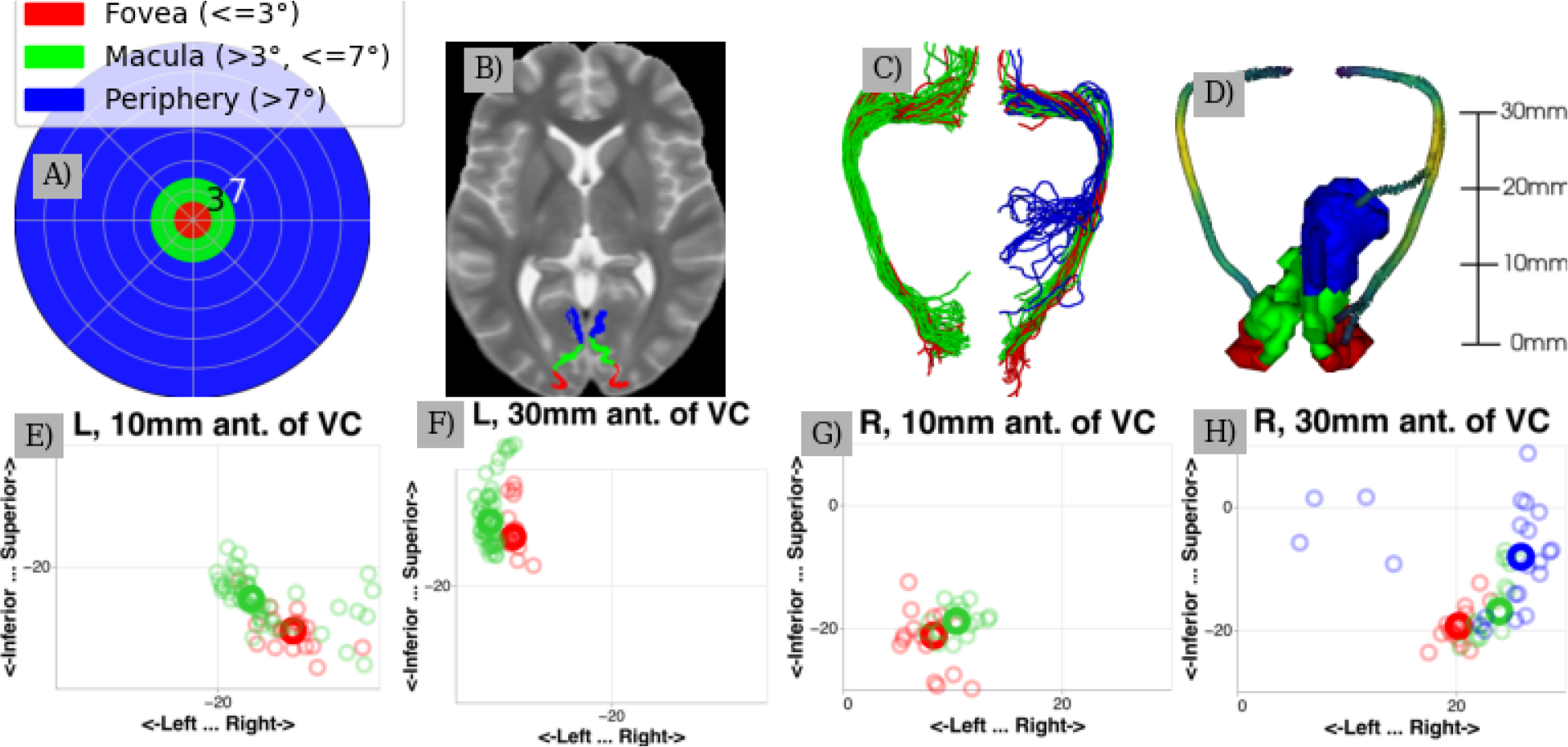
A) Visual field colored by sub-bundle divisions. B) Sub-bundle divisions shown in V1 over the Montreal Neurological Institute (MNI) T2 template ^26^. C) The streamlines identified in an example subject, colored by sub-bundle. Note the left peripheral sub bundle is not found in this subject. D) The core bundles used to extract tract profiles in the example subject. E-H) Slices of the coronal plane along the OR in the example subject. Smaller circles indicate where streamlines pass through the slice, while larger bolded circles indicate where the core bundle passes through the slice. Note that, although the distance between the core bundles may become small (particularly between the macular and foveal sub-bundles), separation between the sub-bundles is maintained along the bundle.

#### Tract profiles

The dMRI signal in each voxel was modeled using the diffusional kurtosis model, implemented in DIPY ^17,31^. The streamlines in each bundle were resampled to 100 points and tissue properties were referred to points along the length of fOR/pOR by extracting the values from the voxels in which each node of each resampled streamline was positioned. Contributions from each node were inversely weighted by their distance from the *core fiber*, the median of the coordinates in each of the 100 nodes along the length of the bundle. In visualizing the results and in statistical analysis, we excluded 20 nodes from either side of the bundle, where tissue properties reflect partial volume effects with the gray matter. An adjusted contrast index (ACI), interpretable as percent difference, is calculated at each position as 2(x_1_-x_2_)/(x_1_+x_2_) for tissue properties (FA, MD, or MK) x_1_ and x_2_ from two bundles. The ACI is used to assess differences along a profile. ACIs can be calculated between sub-bundles or across hemispheres. After calculating ACI and tract profiles for each subject, we display mean profiles / ACI with 95% confidence intervals, calculated using bootstrapping across subjects with 10,000 resamples.

#### Modeling aging

Change over time was assessed using linear regression of the metrics for the mean of nodes from each sub-bundle of the OR with subjects’ age. Errors were estimated using profile likelihood confidence intervals. Additionally, we calculate the mean tissue property (FA, MD, MK) across all nodes for subjects in the youngest age bin. For each tissue property, we used analysis of variance (ANOVA) to quantify the effects of various predictors on variation in the mean tissue property. These predictors are: age, subbundle, hemisphere, and the interaction between age and subbundle.

#### Control bundles

We performed the same analysis on two control bundles, the corticospinal tract (CST) and uncinate (UNC). These bundles were not further divided into subbundles. We used the default ROIs for these bundles as provided in pyAFQ ^1^.

#### Software

All code to reproduce the analysis and the figures is available at <link to be added upon article acceptance>.

## Results

We delineated the trajectory of the fOR and pOR in a sample of 6,021 subjects from the UK Biobank dataset between the ages of 45 and 81. In a portion of these individuals, we were not able to delineate some of the sub-bundles: left fOR in 20.3% of subjects, right fOR in 19.2%, left mOR in 15.7% of subjects, right mOR in 19.8%, left pOR in 22.4%, and right pOR in 34.4%. Successfully delineated tract profiles and their ACIs are visualized (Figs. 3, 4, and 5).

**Figure 3:**
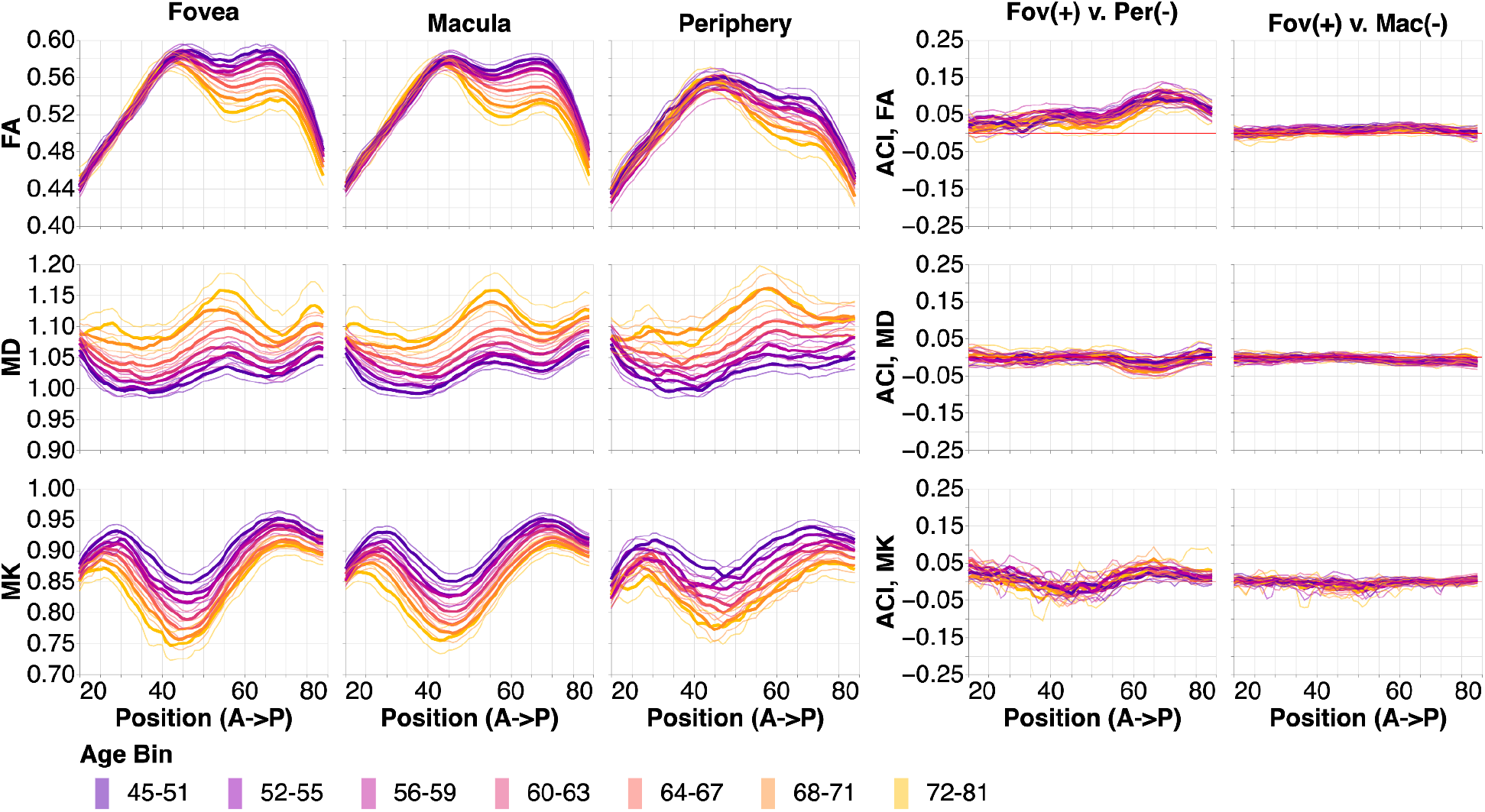
Tissue property profiles along the foveal, macular, and peripheral OR (fOR, mOR, pOR) in the left hemisphere. Positions are from anterior (A) to posterior (P). Subjects are broken down into 7 age bins. The first and last age bin have a larger range of ages so that the number of subjects in each age bin are within the same order of magnitude. In the left two columns, tissue properties are plotted by age bins (different line colors: purple is youngest and gold is oldest). Older subjects tend to have lower FA, higher MD, and lower MK. The thin lines show bootstrapped 95% confidence intervals and are also colored according to age bin. In the right column, we show the adjusted contrast index (ACI) between sub-ORs. Here, higher ACI corresponds to higher values in the tissue properties in the fOR than the pOR or mOR. These differences change only slightly with age, and differences are more pronounced in the posterior section of the OR.

**Figure 4:**
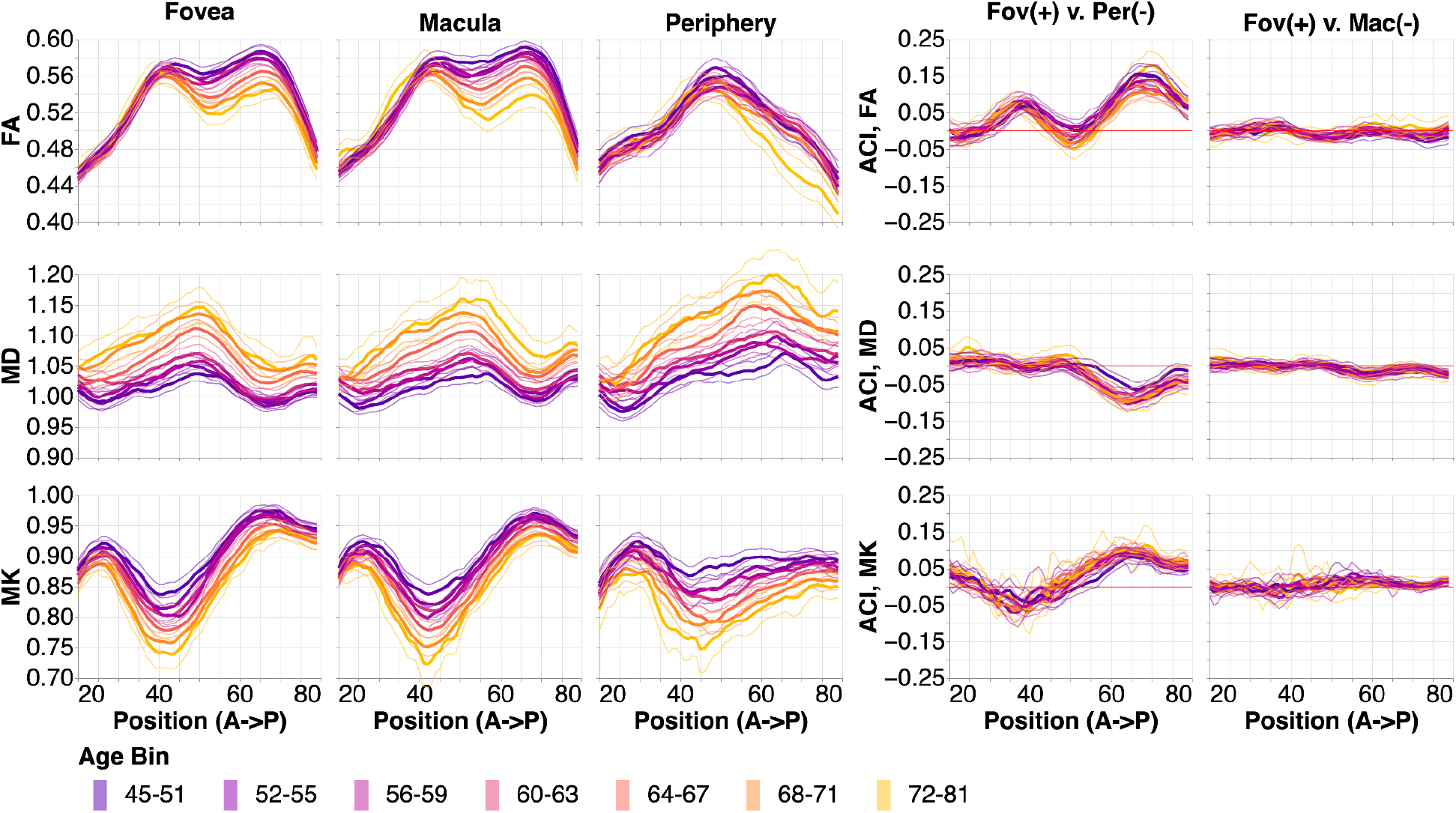
Tissue property profiles along the foveal, macular, and peripheral OR (fOR, mOR, pOR) in the right hemisphere. Positions are from anterior to posterior. Subjects are broken down into 7 age bins. The first and last age bin have a larger range of ages so that the number of subjects in each age bin are within the same order of magnitude (see Figure 1). In the left two columns, tissue properties are plotted by age bins (different line colors: purple is youngest and gold is oldest). The thin lines show bootstrapped 95% confidence intervals and are also colored according to age bin. In the right column, we show the adjusted contrast index (ACI) between sub-ORs. Here, higher ACI corresponds to higher values in the tissue properties in the fOR than the pOR or mOR. These differences change only slightly with age, and differences are more pronounced in the posterior section of the OR.

**Figure 5:**
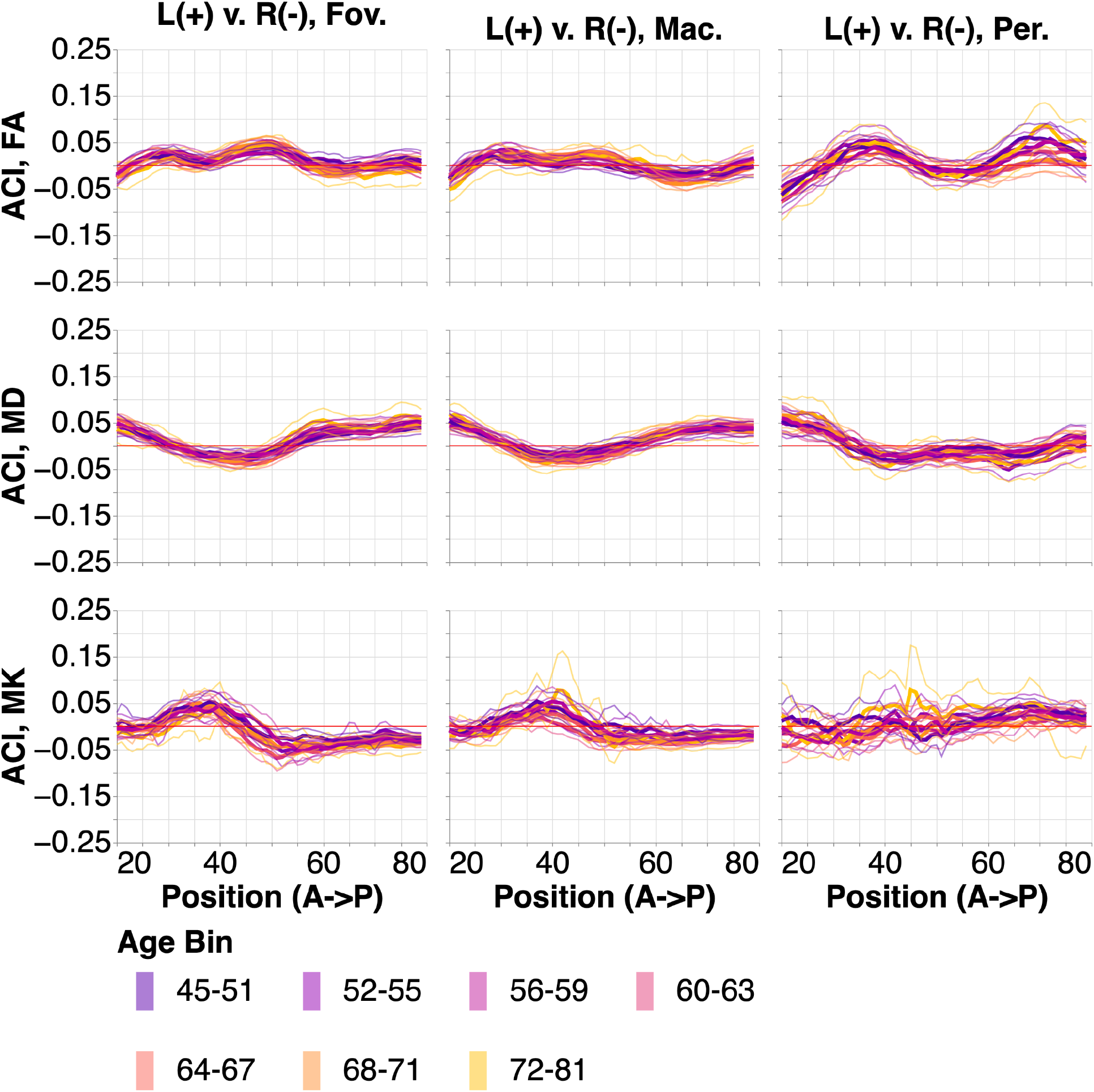
ACI between the left and right ORs. Positions are from anterior to posterior. Subjects are broken down into 7 age bins. The thin lines show bootstrapped 95% confidence intervals and are also colored according to age bin. Here, higher ACI corresponds to higher values in the tissue properties in the left OR than the right OR.

We examine the results in the left hemisphere first (Figure 3). For all tissue properties (Figure 3; rows) and in all sub-bundles (Figure 3; columns), there are consistent changes in with age across almost the entire profile. FA decreases with age, MD increases with age, and MK decreases with age. We conducted pairwise comparisons between points along each of the bundles using a within-subject adjusted contrast index, akin to a percent difference. In FA, there are clear differences between fOR and pOR sub-bundles (the ACI profiles deviate from the red line at 0% difference; Figure 3, under “Fov(+) v. Per(−)”), but it is unclear if there are specific points along the profile where these difference systematically vary with age (different colored lines do not follow a clear gradient). In contrast, there are no points along the profiles, where MK and MD clearly differ between fOR and pOR (the profiles follow the red ACI=0 line closely; Figure 3, under “Fov (+) v. Mac (−)”). Similarly, fOR and mOR sub-bundles do not differ much in any of the tissue properties.

This pattern of results broadly replicates in the right hemisphere (Figure 4). However, in the right hemisphere, there are distinct differences between fOR and pOR in MK and MD in particular points along their profiles that are not apparent in the right hemisphere (Figure 3).

In addition to these differences in the patterns of results between the hemispheres, we also find overall differences between the hemispheres, apparent in point-by-point comparisons of the right and left hemisphere instance of each sub-bundle (Figure 5). Even the consistent point-by-point differences are generally not very large (Almost all ACI are smaller than 5%) and they do not clearly change with age.

We compared the tissue properties in the OR to two non-visual sub-bundles that we used as a point of comparison in a previous study ^14^: the corticospinal tract (CST) and the uncinate fasciculus (UNC). We were not able to delineate left CST in 0.1% of subjects and right CST in 0.1% of subjects. We successfully delineated both UNC in all subjects. Successfully delineated tract profiles and bilateral ACIs are shown in Figures 6 (CST) and 7 (UNC).

**Figure 6:**
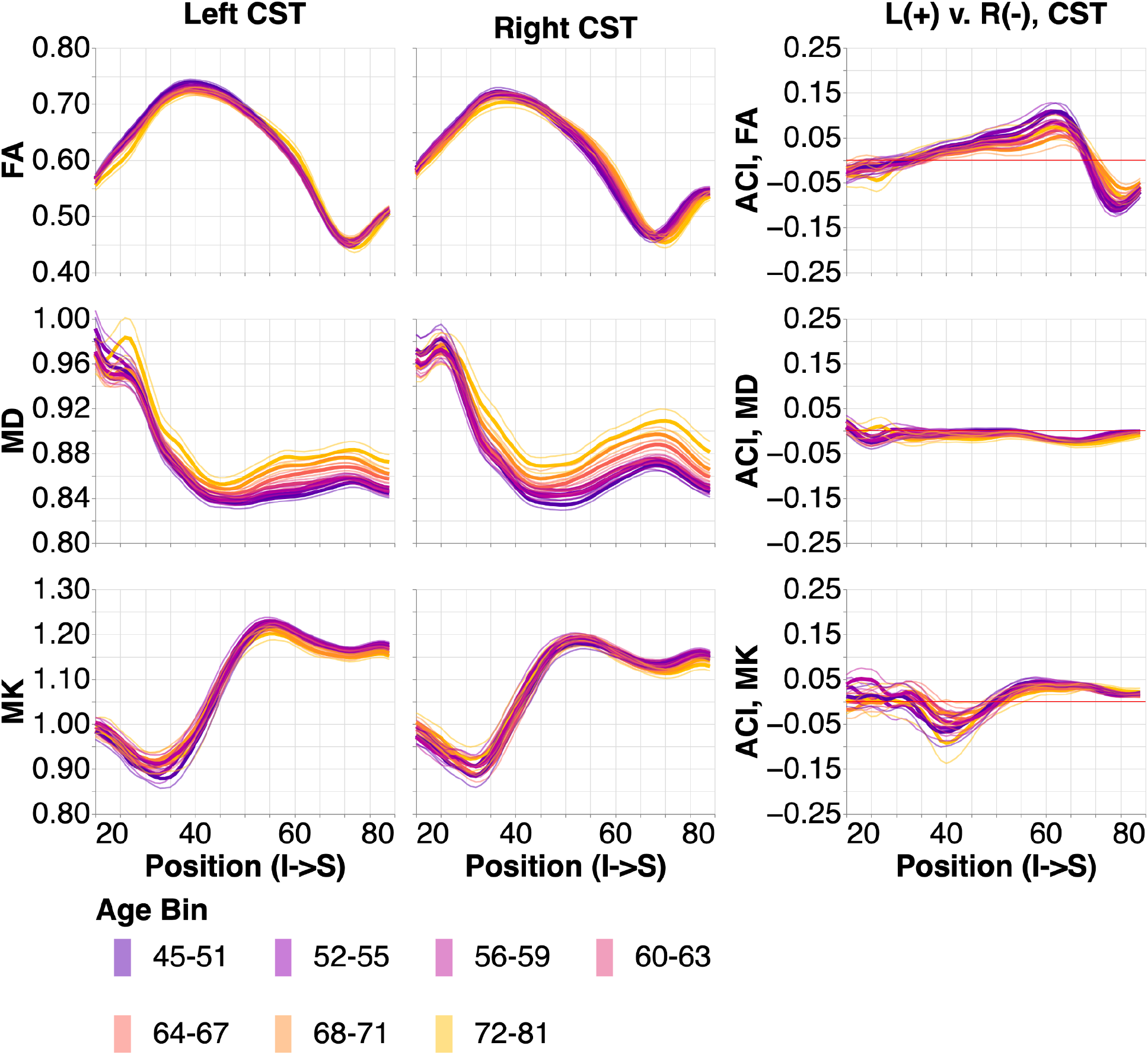
Tissue property profiles along the corticospinal tract (CST). Positions are from inferior to superior. Subjects are broken down into 7 age bins. The first and last age bin have a larger range of ages so that the number of subjects in each age bin are within the same order of magnitude. In the left two columns, tissue properties are plotted by age bins (different line colors: purple is youngest and gold is oldest). The thin lines show bootstrapped 95% confidence intervals and are also colored according to age bin. In the right column, we show the adjusted contrast index (ACI) between the left and right CST. Here, higher ACI corresponds to higher values in the tissue properties in the left CST than the right CST.

**Figure 7:**
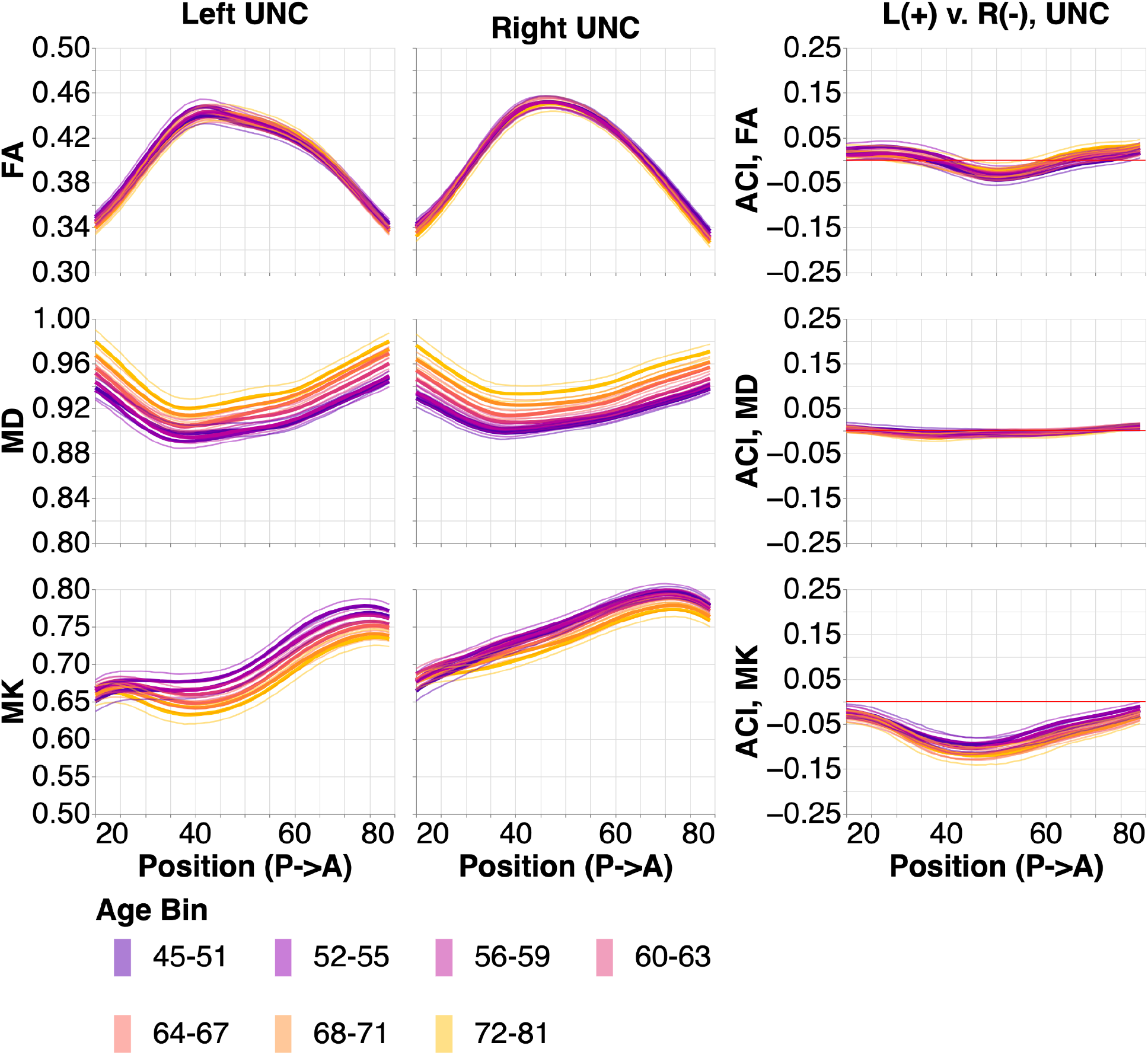
Tissue property profiles along the uncinate (UNC). Positions are from posterior to anterior. Subjects are broken down into 7 age bins. The first and last age bin have a larger range of ages so that the number of subjects in each age bin are within the same order of magnitude. In the left two columns, tissue properties are plotted by age bins (different line colors: purple is youngest and gold is oldest). The thin lines show bootstrapped 95% confidence intervals and are also colored according to age bin. In the right column, we show the adjusted contrast index (ACI) between the left and right UNC. Here, higher ACI corresponds to higher values in the tissue properties in the left UNC than the right UNC.

In the CST we see consistent tissue property profile changes with age (Figure 6) that are similar in their directions (FA decrease, MD increase, and MK decrease with age), but smaller in their magnitude than the changes with age in the OR sub-bundles. CST also has some systematic left-right asymmetries that are particularly large in FA and small in MD. FA profile asymmetries systematically decrease with age.

In the UNC (Figure 7), we again observed changes with age that qualitatively resemble the changes we observed in the OR and CST. These changes were smaller than the changes with age in the OR sub-bundles but larger than the changes observed in CST. Hemispheric asymmetries along the length of the UNC are particularly large in MK and these MK asymmetries increased with age.

To further analyze the tissue properties in the bundles, we averaged each of these quantities along the length of the profiles for every subject and every bundle/sub-bundle. Analysis of tissue profile means recapitulated some of the results that were observed in the point-by-point analysis, and revealed some new observations. For example, in this analysis, we can see that in the CST, mean FA is higher, mean MD is lower, and mean MK is higher than in both the OR sub-bundles. In UNC, mean FA, mean MD, andmean MK are lower than in the OR sub-bundles. OR sub-bundles are similar to each other, but the peripheral OR tends to have lower FA, higher MD, and lower MK.

Using an ANOVA, we model the averaged tissue properties in the OR sub-bundles in terms of hemisphere, aging, and sub-bundle. As expected from the point-by-point analysis (Figures 3,4), we found that mean FA significantly decreases with age (F_1,5530_=237.2, p<0.00001). Even while accounting for age, there are also significant differences between the sub-bundles representing different parts of the visual field (F_2,11060_=4139.3, p<0.00001), presumably because of the lower mean FA in the pOR relative to fOR and mOR (Figure 8). In the aggregate, the small differences seen in the point-by-point hemispheric asymmetry analysis do also constitute a significant lateralization effect, with a higher mean FA in the left hemisphere than in the right hemisphere (F_1,5530_=215.9, p<0.00001). In addition, the effects of age on FA did not affect all sub-bundles uniformly, indicated through a significant interaction between age and subbundle (F_2,11060_=46.2, p<0.00001). We explore this effect in more detail below. MD significantly increases with age (F_1,5530_=701.6, p<0.00001). We also found a lateralization effect in MD, with higher MD in the left hemisphere than in the right (F_1ι5530_=68.8, p<0.00001), and a sub-bundle effect (F_2,11060_=608.0, p<0.00001). Here, interaction between sub-bundle and age is not significant. MK significantly decreases with age (F_1,5530_=587.9, p<0.00001), is significantly higher in the right than in the left hemisphere (F_1,5530_=276.6, p<0.00001), and has a sub-bundle effect (F_2,11060_=709.7, p<0.00001). Again, interaction between sub-bundle and age is not significant for MK.

**Figure 8:**
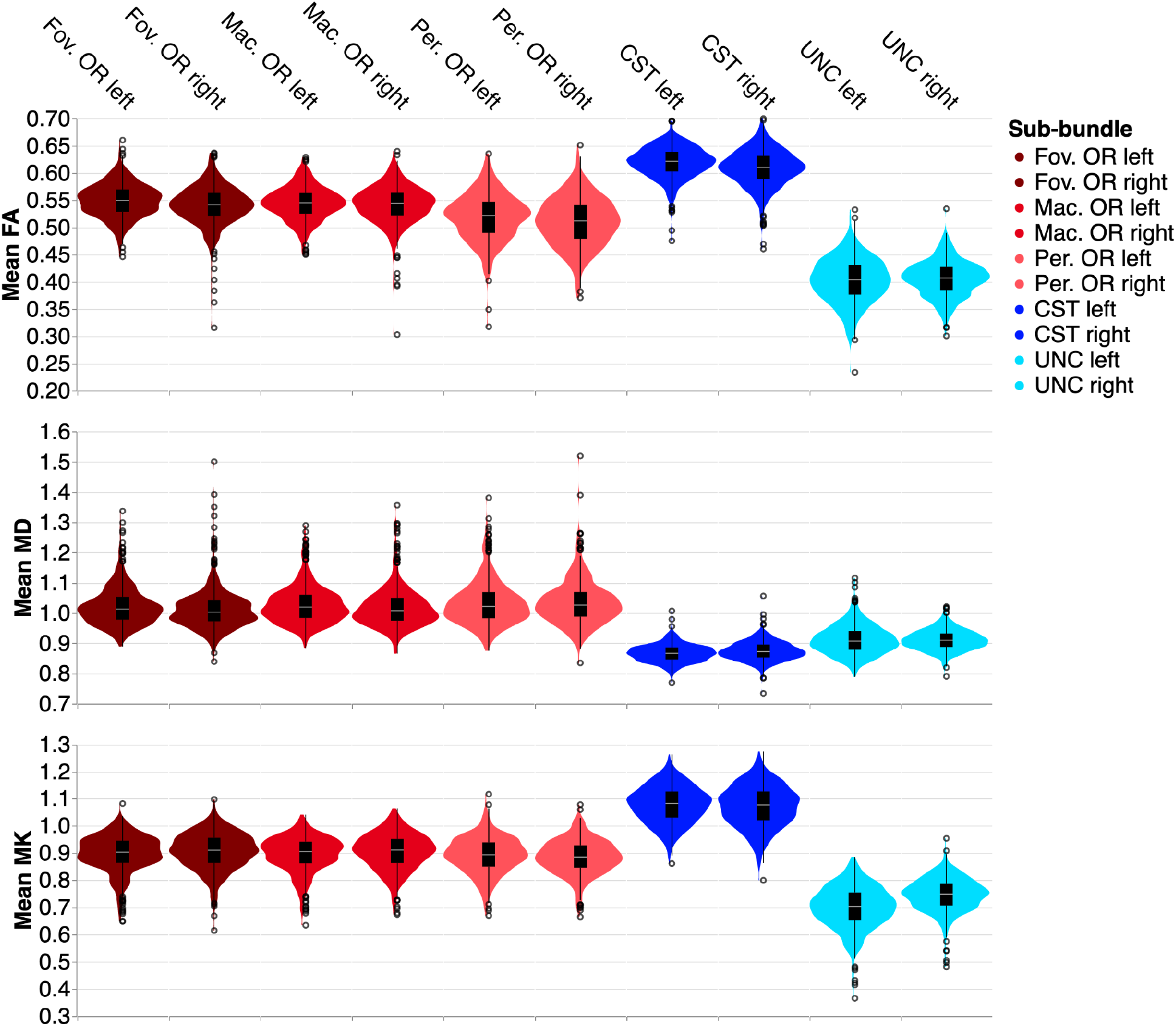
Distribution of mean microstructural tissue properties in the youngest age group (45-51). Note that pOR subbundles have slightly lower mean FA, higher mean MD, and lower mean MK.

To further understand the manner in which age affects the averaged tissue properties, we fit a separate linear regression model to the mean of each tissue property in each bundle sub-bundle (Figure 9). This allowed us to quantify the average rate of change in a tissue property. OR sub-bundles all change substantially more rapidly with age than the control bundles, in all three tissue properties, indicated by linear regression slopes of larger magnitudes. The significant age by sub-bundle ANOVA interaction is explained by the consistent differences between the rate of change in the two central visual field OR sub-bundles and pOR: FA decreases more rapidly in fOR and mOR than in pOR. Much smaller differences in rate of change are observed in MK and MD. However, MD in the pOR increases at a faster rate than in the fOR and mOR.

**Figure 9:**
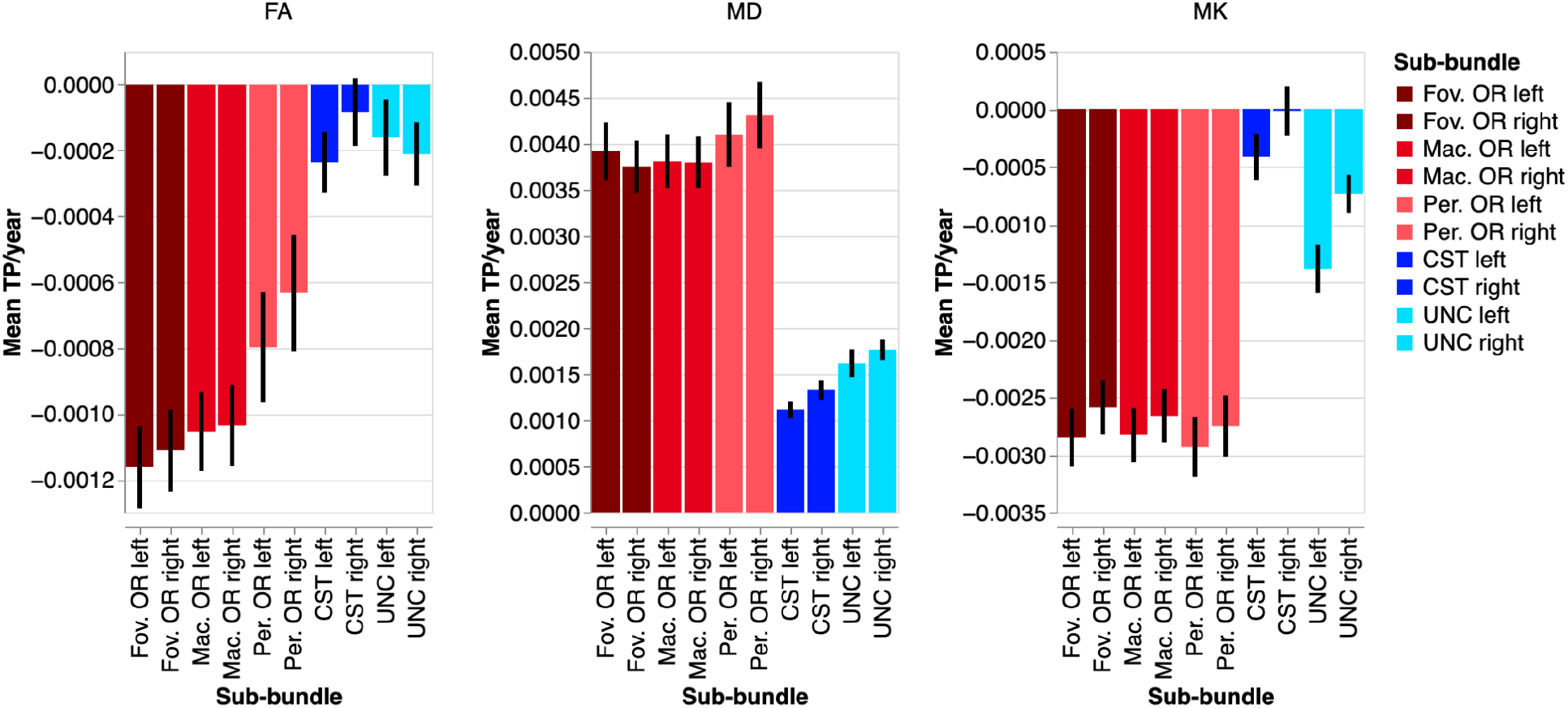
Change in microstructural tissue properties per year according to a linear regression of the mean of each metric. Error bars show the 95% confidence interval. Note that fOR/mOR change more dramatically with age in mean FA than pOR, and that the OR sub-bundles change more with age than the control bundles.

## Discussion

Consistent with results from post-mortem dissections ^32^, we found sub-bundles that follow different anatomical trajectories to different parts of the visual cortex. In addition, we found that the microstructural tissue properties in the pOR differ from those measured in the fOR/mOR. We found higher FA, lower MD and higher MK in fOR/mOR relative to pOR. Using the diffusional kurtosis model (DKI) allows us to disambiguate changes in tissue properties ^16,17^ and taken together, these differences are consistent with more densely packed and coherently-oriented white matter in the foveal/macular OR relative to the peripheral OR. We also found that the mean tissue properties in the OR overall age more rapidly than the tissue properties in two control bundles that we analyzed: the corticospinal tract (CST) and the uncinate fasciculus (UNC). The relatively faster aging in OR is consistent with previous results in the UKBB, with a sample roughly half the size of the current sample ^7^.

Within the OR, we found that all sub-bundles age in a manner that is consistent with age-related declines in density and tissue organization (decreased FA, increased MD and decreased MK). However, concomitant faster declines of FA in fOR/mOR and faster increase in MD in the pOR suggest distinct aging processes happening in parallel. Overall, parts of the visual field with higher-resolution vision (fovea/macula) are associated with white matter bundles that have higher FA, lower MD, and higher MK. Taken together, these two sets of findings are consistent with the high degree of information transmission that needs to be handled by the optic radiations, and particularly within the foveal and macular portions. Replicating previous results ^12,33,34^, we also found consistent lateralization effects, with higher FA, higher MD and lower MK in the left than in the right hemisphere.

The study and our conclusions are still subject to several limitations. First, automatically detecting the OR within every individual is a challenging computational task, particularly across a large and diverse sample (e.g., in terms of their ages). This is because of the high curvature of the tract, its narrow path leading into the occipital pole and its intersection with multiple other pathways, challenges which could be compounded by the expansion of the lateral ventricles with age. Moreover, defining the sub-bundles of the OR is also challenging and we were not able to define the OR bundles or sub-bundles in some of the subjects. In addition to the large variance in subject ages, this could reflect variable data quality among different individuals in the sample, and reflects the difficulties of consistent tractography to cortical (i.e., V1) and small subcortical (i.e., LGN) targets. Still, the advantages of the UKBB dataset are clear: it provides a very large sample, providing high confidence in the consistent results that we see here. Finally, tractography cannot differentiate feedforward axons that transmit information from the LGN to cortex from the feedback projections that transmit information in the opposite direction. Though these feedback projections are thought to be abundant ^35^, their relative volume fraction within the bundle is not well known. Thus, our conclusions need to be viewed as encompassing the properties of both feedforward and feedback projections within the OR.

To summarize, our findings show that the white matter pathways carrying information from different parts of the visual field have distinct biological properties. The largest differences occur between the pOR and fOR/mOR, which follow different anatomical trajectories. The fOR/mOR have properties consistent with the higher degree of information transmission in this part of the visual field and follow a distinct aging process relative to the pOR. Future research could continue to study whether these differences between white matter representing different parts of the visual field is inherited by other structures further into the visual system. For example, the retinotopic organization of the OR is also apparent in the callosal tracts that connect the visual cortex in both hemispheres ^36^ and a similar analysis could be applied to the sub-bundles of the corpus callosum. The methods used here to delineate the different sub-bundles of the OR could be carried forward into population studies of visual disorders that differentially affect different parts of the visual field, such as age-related macular degeneration, as has already been done in small samples ^15^. More generally, the findings demonstrate consistent anatomical variation in tissue properties and their aging even within a single white matter pathway.

## Data Availability Statement

This study uses publicly available data from the UK Biobank. More information on the data and access can be found here: https://www.ukbiobank.ac.uk/enable-your-research

